# PKB/AKT attenuates Lewy Body-like pathology in primary neurons via Cathepsins B and D

**DOI:** 10.1101/2022.09.23.508066

**Authors:** Julia Konovalova, Safak Er, Justyna Barut, Katarzyna Rafa-Zabłocka, Kelvin C. Luk, Małgorzata Figiel, Mikko Airavaara, Andrii Domanskyi, Piotr Chmielarz

**Affiliations:** Institute of Biotechnology, HiLIFE, FI-00014 University of Helsinki, Finland; Drug Research Program, Division of Pharmacology and Pharmacotherapy, Faculty of Pharmacy, FI-00014 University of Helsinki, Finland; Department of Brain Biochemistry, Maj Institute of Pharmacology, Polish Academy of Sciences, Kraków, Smętna 12, Poland; Department of Pathology and Laboratory Medicine, Center for Neurodegenerative Disease Research, Perelman School of Medicine, University of Pennsylvania, Philadelphia, PA 19104; Department of Physical Biochemistry, Faculty of Biochemistry, Biophysics and Biotechnology, Jagiellonian University, Kraków, Poland; Neuroscience Center, HiLIFE, 00014 University of Helsinki, Finland

**Keywords:** AKT, Parkinson’s disease, alpha-synuclein, misfolded protein accumulation, autophagy, cathepsin

## Abstract

Motor symptoms of Parkinson’s disease (PD) are caused by the loss of dopamine neurons in the substantia nigra pars compacta. Current PD therapies offer only symptomatic relief, and the research on novel ways to stop or slow down the degeneration of dopamine neurons in PD attracts considerable efforts.

A characteristic feature of PD is the abnormal accumulation and spread of misfolded α-synuclein (α-syn) protein leading to the formation of Lewy neurites and bodies in different neuronal populations, including dopamine neurons. The process of Lewy body formation compromises neuronal functions and contributes to the progression of the disease and neurodegeneration in PD patients. Treatments preventing α-syn misfolding, aggregation and/or spread may therefore provide much-needed disease-modifying therapies for PD. We and others have demonstrated that α-syn aggregation and Lewy body formation can be modelled in primary neuronal cultures treated with α-syn pre-formed fibrils (PFFs). Using this approach, we have identified several factors preventing the accumulation of phosphorylated α-syn in dopamine neurons *in vitro* and *in vivo*. In particular, we and others have demonstrated a prominent role of PKB/AKT pathway activation in promoting the survival of dopamine neurons.

Here, we utilize lentiviral vectors (LVs) to express constitutively active myristoylated serine-threonine protein kinase AKT1 (mAKT1) in primary dopamine neurons. We show that lentivirally-delivered mAKT1 promoted survival of dopamine neurons after thapsigargin-induced endoplasmic reticulum (ER) stress and prevented fibril-induced accumulation of phosphorylated α-syn. Furthermore, using pharmacological tools, we demonstrate that this activity of mAKT1 is dependent on the functional Cathepsins B and D. These proof-of-principle results demonstrate the critical importance of lysosomal enzymes for processing misfolded α-syn and establish a basis for the use of LVs to develop neuroprotective gene therapy strategies for dopamine neurons in PD.

## Introduction

Synucleinopathies, including Parkinson’s disease (PD), represent a group of diseases characterized by abnormal accumulation of α-synuclein (α-syn) protein [1]. In the brains of both sporadic and familial PD patients, aggregated and mostly Ser129-phosphorylated α-syn (pαSyn) containing inclusions – called Lewy bodies – are observed in different neuronal populations, including dopamine neurons in the substantia nigra and other brainstem nuclei like noradrenergic neurons in locus coeruleus [2,3]. Lewy pathology is also prominent in other brain areas, e.g. in the hippocampus, frontal cortex and amygdala [2]. Furthermore, Lewy pathology is present in enteric neurons and in the peripheral nervous system and might be highly relevant in these areas for disease etiology as proposed in the body-first PD hypothesis [4]. Accumulating pαSyn-positive neuronal aggregates is thought to compromise neuronal functions even at the early stages of PD by causing synaptic and mitochondrial dysfunction, cellular stress, and deregulation of protein degradation pathways [5,6]. Over time, Lewy pathology spreads to the connected areas of the central nervous system, and this spreading may correlate with the disease severity [2].

Activation of AKT/mTOR signalling by either Pten ablation [7,8] or expression of constitutively active myristoylated AKT1 (mAKT1) [9,10] promotes axonal growth and sprouting and protects dopamine neurons in mouse PD models [8,11]. Furthermore, we recently reported that AKT and Src pathways are involved in the protective action of glial cell line-derived neurotrophic factor (GDNF)/RET signalling against α-syn aggregation [12]. In addition, pharmacological inhibition of AKT exaggerates α-syn aggregation in dopamine neurons in a dose-dependent manner. However, the complete mechanism of putative action of AKT on α-syn accumulation is not yet fully understood.

Misfolded α-syn is cleared by both proteasomal and lysosomal protein degradation pathways [13]. In particular, misfolded α-syn species can be degraded by lysosomal cathepsins D, B and L [14–17]. The endo-lysosomal pathway has also been implicated in the intercellular transmission of pathological α-syn. Extracellular α-syn is uptaken via endocytosis and subsequently transported to lysosomes, where it escapes by an unknown mechanism [18]. Uptaken α-syn fibrils may persist in neuronal lysosomes for days before their release to the cytoplasm [19]. During this time, α-syn fibrils are progressively fragmented into smaller species [20]. Therefore, the ability of the cell to degrade uptaken α-syn and to maintain endo-lysosomal integrity might dictate the rate of cell-to-cell transmission of pathological α-syn. Notably, the role of lysosomal function in PD progression is also supported by genetic evidence. Indeed, heterozygous mutations in lysosomal glucocerebrosidase (GBA) are the largest genetic risk factor for developing familial PD [21,22], while variants in lysosomal ion channel TMEM175 have also been strongly implicated [23,24].

In this study, we aimed to confirm the protective role of AKT1 against the accumulation of α-syn aggregates in the neurons and elucidate the underlying mechanisms, particularly possible links to the regulation of the endo-lysosomal pathway. We utilized lentiviral overexpression of constitutively active mAKT1 and pre-formed fibrils (PFFs)-induced model of α-syn aggregation in primary mouse embryonic dopamine and hippocampal neurons *in vitro* [6,25,26] and cortical neurons *in vivo*. Our results show that lentiviral overexpression of mAKT1 promotes the survival of primary dopamine neurons in basal and thapsigargin-induced endoplasmic reticulum (ER) stress conditions. We further demonstrate that mAKT1 significantly attenuated α-syn aggregation in the soma of dopamine and hippocampal neurons. We also confirmed its effectiveness *in vivo* in cortical neurons. Furthermore, mAKT1 overexpression led to significant changes in the expression of autophagy-related genes and lysosomal proteases. Inhibition of these proteases abolished the positive effect of mAKT1, suggesting that autophagy/endo-lysosomal pathway regulation is the plausible mechanism of action for AKT-mediated elimination of α-syn aggregation in neurons.

## Material and Methods

### Construction of transfer plasmids

A 448 nt fragment of human *SYN1* promoter was PCR amplified with Phusion high fidelity DNA polymerase (Thermo Scientific) and the following primers: BcuI_hSYN_for 5’-AT<u>ACTAGT</u>AGTGCAAGTGGGTTTTAGGACC and EcoRI_hSYN_rev 5’-TG<u>GAATTC</u>GACTGCGCTCTCAGG, then cloned into FastDigect-BcuI/-EcoRI (Thermo Scientific) digested pCDH-CMV-MSC-T2A-EGFP vector (System Biosciences). mAKT1 (Merck, #21-151) and human GDNF cDNA [12] were each cloned into FastDigest-EcoRI/-NotI-digested pCDH-hSYN-EGFP vector.

### Production of lentiviral particles

Linear polyethylenimine (PEI, MW 25000; Polysciences Inc, #23966-2) was dissolved in 1× PBS (pH 4.5) by heating for 30-40 min in a 75°C water bath, then cooled to room temperature (RT) and sterilized by filtering through a 0.22 μm syringe filter (GE Healthcare Life Sciences, WHA10462200); aliquots were stored at 4°C. To produce lentiviral vectors, ≈ 3×10^6^ human embryonic kidney 293T cells (ATCC, #CRL-1573) were plated on 10-cm cell culture dishes in Dulbecco’s Modified Eagle Medium (DMEM) (Sigma, D-7777) supplemented with 10% fetal bovine serum (Gibco, 10500056), 100 μg/ml Normocin (Invivogen, #ant-nr-2) and 25 mM HEPES (VWR Chemicals, 441487M). After 24h, cells were transfected with 4 μg of corresponding transfer plasmid and 2 μg each of the four helper plasmids: pMDLg/pRRE (Addgene, #12251), pRSV/REV (Addgene, #12253), pMD2.G (Addgene, #12259) and pAdvantage (Promega, E1711, not used for LV-GDNF production). The plasmids were mixed with 100 μl of PEI (4:1 v/w PEI:DNA ratio) in 500 μl of pre-warmed OptiMEM II reduced serum medium (Gibco, 1985070), after 10-15 min incubation at RT, 600 μl of transfection mix was added dropwise to cells. After ∼72 h, the supernatants were pooled, centrifuged at 300× *g* for 5 min, filtered through a 0.22-μm pore size filter, and transferred to a centrifuge tube (Beckman UltraClear, #344058) fitted into a metal rotor tube (Beckman AH-629 rotor). The lentiviral particles were pelleted by centrifugation at 120000× *g* for 1.5 h at 4°C, and the pellet was re-suspended in Dulbecco’s phosphate buffered saline, aliquoted and stored at -80°C [12].

### Primary embryonic cultures

All animal experiments were approved by the Finnish National Board of Animal Experiments (license number: ESAVI/7812/04.10.07/2015). NMRI mice (Charles River) were housed in a 12-hour light-dark cycle with free access to food and water. Animals were mated, and E0.5 was defined as the morning when the vaginal plug was detected. Ventral midbrain floors were isolated from E13.5 mouse embryos to establish midbrain cultures, as described previously [25,27]. Cells were plated on 96-well plate in micro islands (35,000 cells/well) and maintained with dopamine neuron medium (DPM) containing 11.5 mM D-glucose (Sigma, G8769-100ML), 2 mM L-glutamine (Gibco, #25030–032), 1× N2 (Gibco, 17502001), 100 μg/mL Primocin (Invivogen; ant-pm) in DMEM/F12 (Gibco, #21331046). To establish hippocampal cultures, hippocampi were isolated from E15.5-E16.5 mouse embryos, as described elsewhere [28]. Cells were plated on 96-well plate (25,000cells/well) and maintained with neurobasal medium (NB) (Life Technologies, 21103-049) containing 1× B-27 Supplement (Gibco, #17504-044), 500 μM L-Glutamine (Gibco, #25030–032), and 100 μg/mL Primocin (Invivogen; ant-pm). All cells were maintained at 5% CO_2_ at 37°C.

Lentiviral particles were diluted in neuronal culture medium (DPM or NB, depending on cultures); 10 μl of diluted lentiviral particles per well were used to transduce the cells at a multiplicity of infection (MOI) ≈5.

Cells were treated with sonicated recombinant mouse α-syn PFFs (final concentration 2.5 μg/ml; more details below) on DIV8 with PBS used as negative control according to a previously established protocol [12,25].

For the induction of ER stress, the midbrain cultures were treated with thapsigargin (Invitrogen, T7458) or vehicle control for 48h. Hippocampal neurons were treated with Bafilomycin A (Sigma, B1793), Lactacystin (AG Scientific, #L-1147), CA-074 (Tocris Bioscience, #4863), SID26681509 (AdooQ BioScience, #A15313), Pepstatin A (Santa Cruz Biotechnology, sc-45036) or corresponding vehicle for 72h. After the treatments, the cells were fixed with 4% paraformaldehyde (PFA) (Aldrich, 166005) for 20 min at RT and immunostained as described below.

### Immunofluorescent staining and quantification of neurons with or without α-syn aggregates

After 4% PFA fixation, cells were washed twice with 1xPBS, and permeabilized for 15 min with PBS containing 0.2% Triton X-100 (Sigma, T9284/X100) (PBST) solution. After blocking unspecific antibody binding sites by 1 h RT incubation in blocking solution (PBST containing 5% normal horse serum (Vector/BioNordika, S-2000)), the cells were incubated with the corresponding primary antibodies diluted in blocking solution: anti-tyrosine hydroxylase (TH) (Millipore, #MAB318; 1:2000) or anti-NeuN (Millipore, MAB377; 1:500) and anti-phosphoSer129-α-synuclein (Abcam, ab51253; 1:2000) overnight at 4ºC. The next day, cells were washed three times with 1xPBS, and incubated with secondary antibodies (goat anti-mouse AlexaFluor 568 (Thermo Fisher Scientific, #A11004) and donkey anti-rabbit AlexaFluor 647 (Thermo Fisher Scientific, #A31573); each 1:500 in PBST) for 1 h at RT. After three washes with 1xPBS, the cells were stained with 4’,6-diamidino-2-phenylindole (DAPI, 0.2 μg/ml in PBS) for 10 min and kept in PBS at 4°C, protected from light until imaging.

The plates were imaged with CellInsight CX5 High Content Screening (HCS) Platform (Thermo Fisher) or ImageXpress Nano Automated Imaging System (Molecular Devices) at 10x magnification, and the images were quantified with CellProfiler and CellAnalyst software packages [25,29,30].

### Western blot

To detect mAKT1 and total α-syn expression, hippocampal cells were seeded on 6-well plates (2*10^6^ cells/well). After treatment with lentiviral particles, cells were lysed in Western Blot (WB) lysis buffer (1% SDS, 10 mM Tris, 1 mM EDTA, phosphatase and protease inhibitors (Roche, 04906837001 and 04693159001, respectively)). Lysates were run on a pre-cast gel (BioRad, 456-1093) at 40 mA for ∼1 h and then blotted to the nitrocellulose membrane (Cytiva, 10600003). The membrane was fixed with 4% PFA + 0.02% glutaraldehyde (Fluka, 49631) for 20 min, incubated in blocking solution (3% skimmed milk or 5% BSA (Sigma-Aldrich, A9647) in TBS + 0.01% Tween20 (Sigma, P2287)) for 1 h at RT and incubated with primary antibodies overnight at +4°C. The membrane was washed and incubated with HRP-conjugated secondary antibodies for 1 h at RT. Signal was detected with Pierce™ ECL Western blotting substrate (ThermoFisher Scientific, 32106) according to manufacturer’s instructions. To detect total α-syn, anti-α-synuclein (Abcam, #ab1903, 1:2000 in 3% milk) and goat-anti-mouse IG (Dako, P0447, 1:3000 in 3% milk) antibodies were used. For AKT1 detection, anti-AKT (pan) (11E7) (Cell Signalling Technology, #4685, 1:1000 in 5% BSA) and donkey-anti-rabbit IG (GE Healthcare, NA9340V, 1:3000 in 5% BSA) antibodies were used. GAPDH (Millipore, MAB374) was used as the housekeeping gene.

### qPCR

Total RNA from hippocampal neurons was isolated using Cell-to-Ct (ThermoFisher Scientific, AM1728) or miRNeasy Micro (Qiagen, 217084) kit according to manufacturer’s instruction and used as a template for cDNA synthesis with Maxima H Minus Reverse Transcriptase kit (ThermoFisher Scientific, #EP0753). Quantitative PCR was performed with LightCycler® 480 System (Roche Molecular Diagnostics) using the LightCycler® 480 Software release 1.5.1.62, using TaqMan Gene Expression Assays (ThermoFisher Scientific) (probe numbers: Mm01188700_m1, Mm00547102_m1, Mm00495262_m1, Mm00448968_m1, Mm01265461_m1, Mm01310506_m1, Mm00515586_m1) according to the manufacturer’s protocol. Expression of genes of interest was normalized to housekeeping genes *Hprt1* (Mm01545399_m1) and *B2M* (Mm00437762_m1) to ensure equal input of cDNA. Each data point represents an average of 2 wells (technical replicates) from 3-4 different experimental plates (biological replicates).

### α-syn fibrils preparation

For *in vitro* induction of α-syn aggregation, α-syn PFFs were prepared by sonicating mouse recombinant α-syn fibrils [31]. PFFs were diluted in 1× PBS to a final concentration of 100 μL/Ml and sonicated with Bioruptor sonication device (Diagenode, Liege, Belgium, #B01020001) at the following settings: 10 cycles, 30 s on/ 30 s off. PFFs were added to the cell culture medium at a final concentration of 2.5 μg/mL. For *in vivo* administration, mouse α-syn monomer was purified employing a modified protocol of Volpicelli-Daley et al. (2014)[31], from BL21 RIL cells transformed with pD454-SR-mα-syn plasmid (Addgene, #89075) and induced with 1 mM IPTG. The cells were disrupted by sonication, and the lysate was precleared by boiling. α-syn was enriched by ion-exchange chromatography with the use of Fractogel-TMAE (Merck) and MonoQ (Cytiva) columns and finally purified by size-exclusion chromatography with a Superdex 75 (GE Healthcare) equilibrated with 10 mM Tris-HCl pH 7.4, 10 mM Na_2_HPO_4_, 100 mM NaCl. Spin-concentrated α-syn monomers were aggregated under shaking for 7 days at 37°C according to described protocols [32]. The obtained fibrils were diluted in 1×PBS to a final concentration of 2 mg/mL and sonicated with a high-power probe sonicator (UP100H, Hielscher) with the following settings: 2 mm sonicator probe, 60 s with maximum power pulses, 0.5 s on/0.5 s off on ice.

### Animals

The experiments were performed on adult C57BL/6J male mice (12 weeks-old) housed at standard conditions at room temperature 22±2°C, 55±10% humidity and 12 h light/dark cycle, with food and water ad libitum. All animal experiments were approved by the Local Ethical Commission for Animal Experiments at Maj Institute of Pharmacology, Polish Academy of Sciences in Krakow (permit number 304/2020) and fulfilled the requirements of the EU Directive 2010/63/EU on the protection of animals used for scientific purposes.

### Stereotaxic surgeries

Stereotaxic injections were performed under ketamine (80 mg/kg) / xylasine (10 mg/kg) anesthesia. Animals were injected with lentiviral vectors (LV-mAKT1-GFP or LV-GFP) unilaterally at coordinates A/P +2.1 mm, M/L +0.3 mm, D/V-2.0 mm from bregma, 1.5 μL at 0.25 μL/min, after the injection the needle was allowed to sit in place for another 3 minutes. One week after lentiviral injection, animals were injected with α-syn PFFs at coordinates A/P 0.7 mm, M/L -2.2 mm, D/V -3.0, 2.5 μL at 0.2 μL/min, after the injection, the needle was allowed to sit in place for another 3 minutes.

### Tissue processing and staining and quantification

30 days after PFFs injection, animals were sacrificed, their brains were extracted and fixed in 4% paraformaldehyde (PFA) for 48 h, and stored in 0.4% PFA. The brains were cut on a vibratome (Leica, Germany) into 40 μm sections. Sections from the frontal cortex were stained with chicken anti-GFP (ThermoFisher, A10262; 1:500) and rabbit anti-pSer129α-syn (Abcam, ab51253; 1:4000) antibodies and goat anti-chicken Alexa488 (ThermoFisher, A11039; 1:500) and donkey anti-rabbit Alexa594 (ThermoFisher, A21207; 1:500) secondary antibodies, mounted with Vectashield (Vector, H-1500) and imaged on wide field mode on Leica TCS SP8 WLL. For quantification, sections from corresponding cortical areas showing successful lentiviral transduction of neurons in cortical layers were chosen from 3 animals from each group. The total number of GFP labeled cells (transduced cells), and the number of GFP labeled cells harboring cytoplasmic α-syn inclusions were quantified, and the percentage of transduced cells harboring α-syn was calculated.

### Statistical analysis

Statistical significance for *in vitro* experiments was calculated by randomized block one-way or two-way repeated measurements ANOVA or mixed-model analysis (Lew, 2007), followed by Dunnett’s or Holm-Sidak multiple comparison tests; for *in vitro* inhibitor data post-hoc analyses were done with Holm-Sidak test separately for DMSO vs inhibitor concentrations and GFP vs mAKT1 for each inhibitor concentration; for *in vivo* data statistical significance was calculated using students t-test. Calculations were done utilizing GraphPad Prism 9.1.0 software (GraphPad Software, Inc., La Jolla, CA, USA). Data are represented as means ± SD.

## Results

### Lentiviral delivery of mAKT1 promotes survival and stress resistance of dopamine neurons

To study the effect of AKT1, a lentivirus vector (LV) expressing myristoylated AKT1 (LV-mAKT1) under the control of human *SYN1* promoter (Figure 1A) was produced. Addition of the myristoylation signal targets AKT1 to the cellular membrane, causing constitutive activation of PKB/AKT pathway [9]. To test the functionality of LV-mAKT1, its ability to promote survival of primary embryonic dopamine neurons was compared to LV expressing only GFP (LV-GFP) as a negative control and LV driving human GDNF expression (LV-GDNF) as a positive control. Transduction of primary midbrain cultures with LV-mAKT1 and LV-GDNF significantly increased the survival of TH-positive dopamine neurons compared to cultures treated with LV-GFP (Figure 1B).

**Figure 1.**
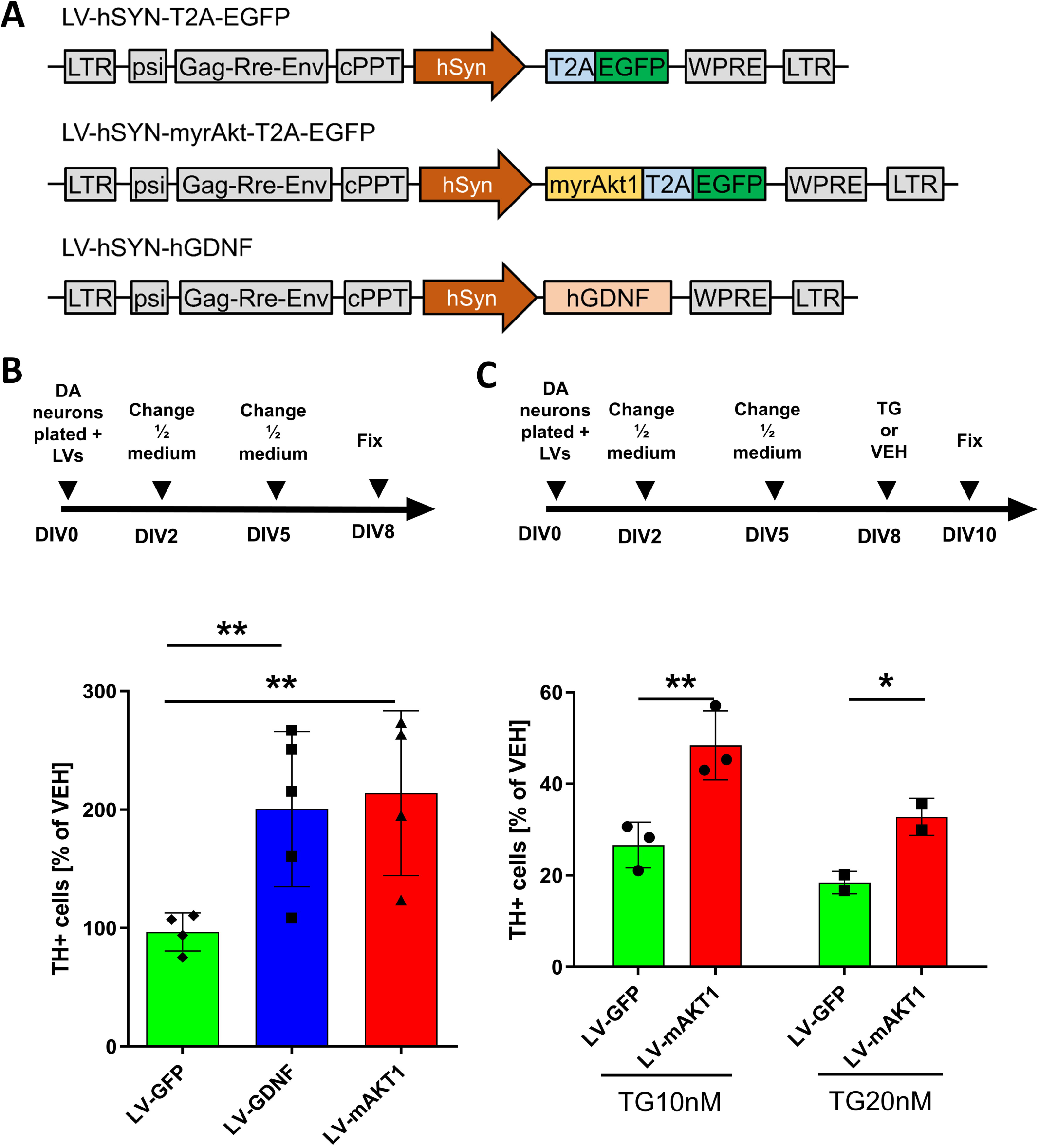
Lentiviral (LV) delivery of myristoylated AKT1 (LV-mAKT1) promotes survival and ER stress resistance of cultured dopamine neurons. A) Maps of plasmid constructs used for the production of LVs to express GFP (negative control), mAKT1 or hGDNF (positive control) in neuronal cultures. B) LV-driven expression of mAKT1 increases survival of TH-positive (TH+) neurons in midbrain primary cultures. C) Overexpression of mAKT1 in midbrain primary cultures protects against ER stress-related cell death induced by thapsigargin. *—p-value < 0.05, **—p-value < 0.01. Data are represented as mean ± SD. Each dot represents a mean value from all technical replicates (wells) in the independent experiment.

In addition, we evaluated the protective effect of LV-mAKT1 on the survival of dopamine neurons under ER stress. To induce ER stress in dopamine neurons, cultures were treated with thapsigargin – an inhibitor of the sarco/endoplasmic reticulum Ca^2+^ ATPase [33]. In accordance with previous findings [34], thapsigargin treatment caused significant loss of TH-positive cells in primary midbrain cultures, while transduction with LV-mAKT1 strongly attenuated the negative effect of thapsigargin treatment (Figure 1C).

### mAKT1 reduces aggregation of Ser129-phosphorylated α-syn in primary dopamine and hippocampal neurons

We have previously demonstrated that AKT is a critical player in the attenuation of α-syn aggregation, and inhibition of AKT exacerbates α-syn pathology in primary midbrain cultures [12]. To further study the effect of AKT on α-syn accumulation, we tested the ability of mAKT1 overexpression to attenuate α-syn pathology in primary dopamine and hippocampal cultures. To do so, a model of α-syn pathology from [12] was used with small modifications. The cells were pre-treated with LVs at DIV5, and PFFs were added 3 days later at DIV8 when LV-mediated GFP expression was observed. Cells were fixed at DIV15 (Figure 2A) when large α-syn aggregates in the cell body were detected. Overexpression of mAKT1 resulted in a significant reduction of α-syn aggregates in both dopamine (Figure 2B) and hippocampal (Figure 2C) neurons, with a more profound effect in the latter one. Neither LV-GFP nor LV-mAKT1 treatment affected neuronal survival in this period (Figure S1A and B).

**Figure 2.**
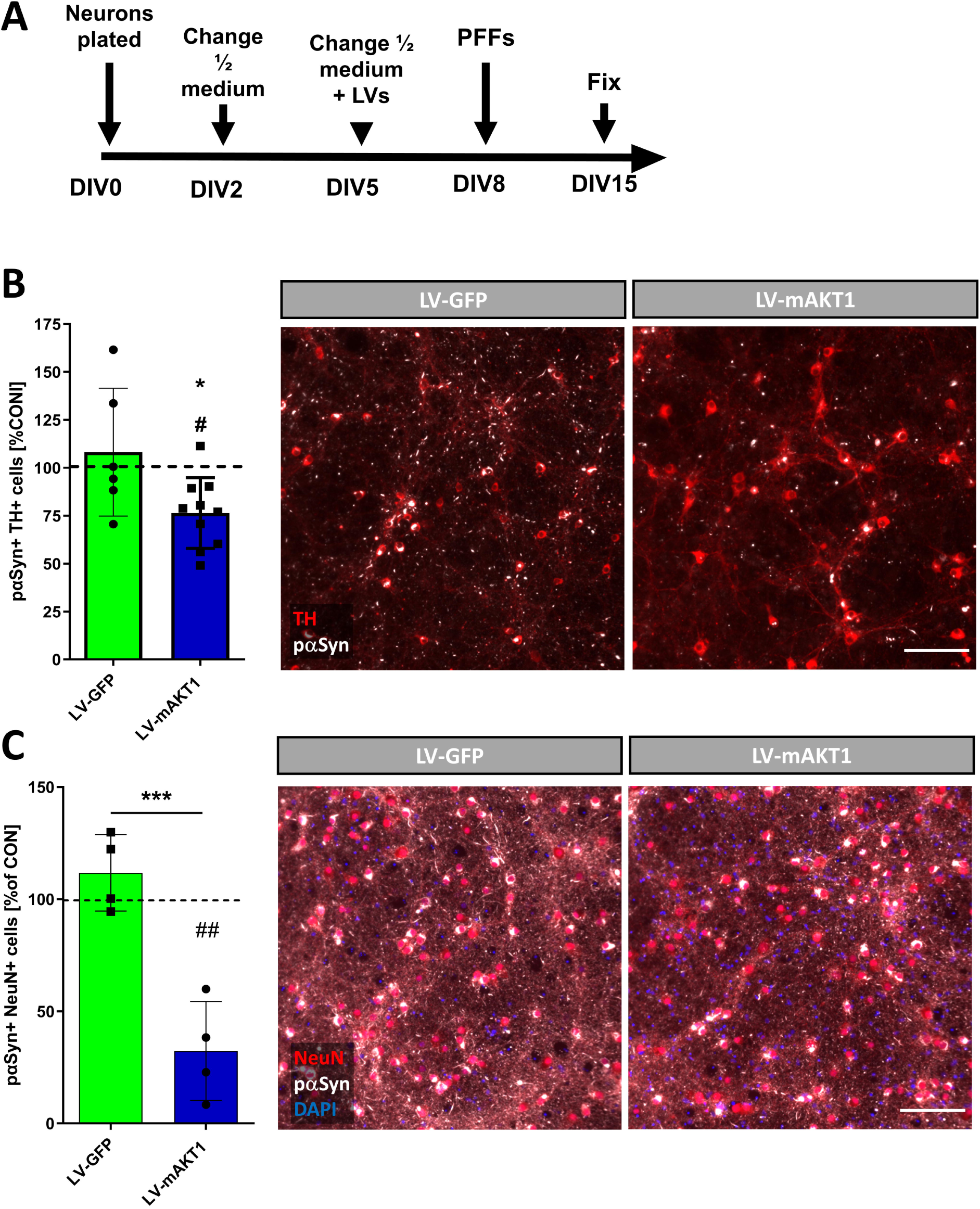
Expression of mAKT1 alleviates α-syn aggregation in primary dopamine and hippocampal neurons. A) Timeline of experiments. B) LV-driven expression of mAKT1 reduces the number of Ser129-phosphorylated α-syn (pαSyn) aggregates in the TH-positive (TH+) neurons of midbrain cultures. C) Overexpression of mAKT1 attenuate the accumulation of α-syn in primary hippocampal neurons (NeuN+). ^*^—p-value < 0.05, ^***^—p-value < 0.001 against LV-GFP, # p<0.05, ## p<0.01 vs PBS control. Data are represented as mean ± SD. Each dot represents a mean value from all technical replicates (wells) in the independent experiment. Scale bars, 400 μm.

### mAKT1 overexpression does not affect expression levels of endogenous α-syn

As it was shown previously, PFFs induce the formation of α-syn inclusions via recruiting endogenous α-syn; importantly, high levels of endogenous α-syn are critical for forming of these aggregates [31]. Therefore, we assessed if mAKT1 overexpression reduces the formation of aggregates through modulation of endogenous α-syn expression. However, no differences were found in α-syn levels measured at mRNA (Figure 3A) and total protein levels (Figure 3B) after mAKT1 overexpression.

**Figure 3.**
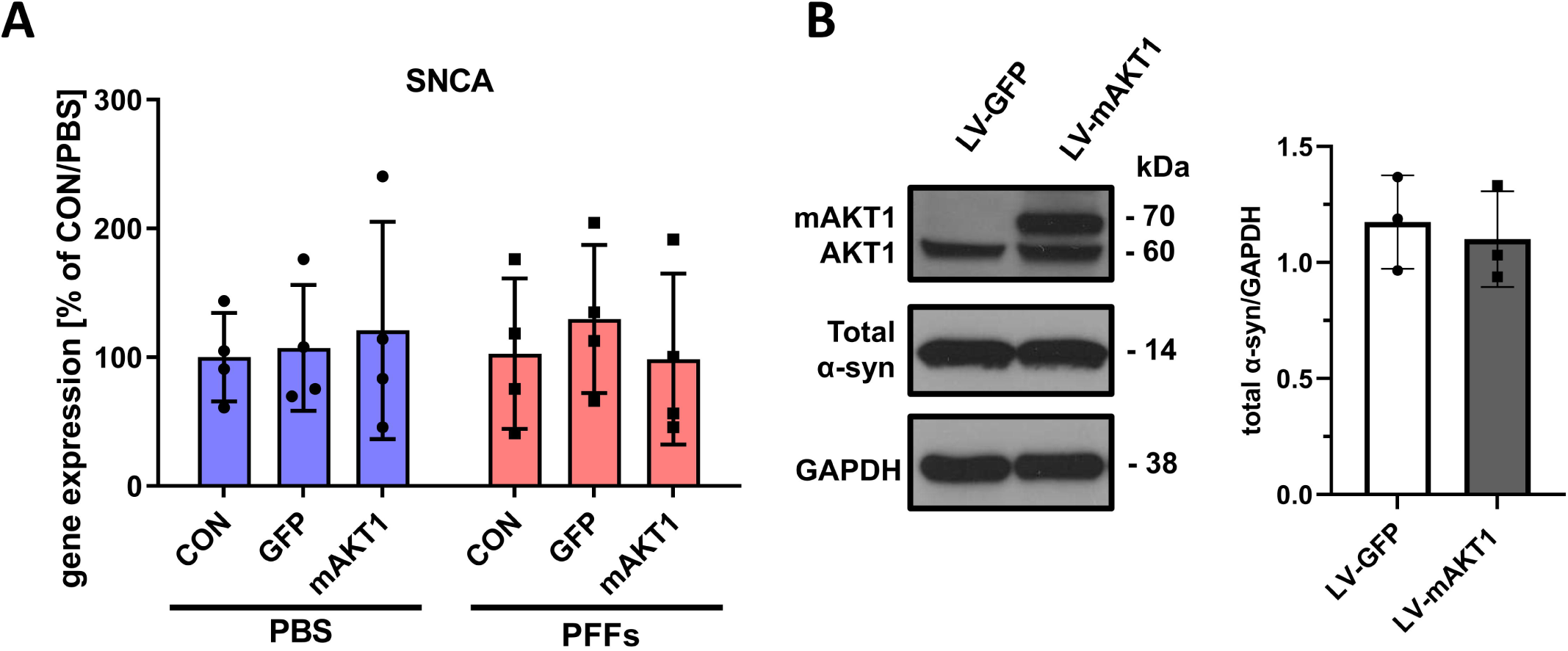
mAKT1 does not affect expression levels of endogenous α-syn. A) Neither LVs nor PFFs treatment had a significant effect on mRNA levels of α-syn (SNCA). B) Protein levels of total α-syn were not changed by mAKT1 expression. Data are represented as mean ± SD. Each dot represents a mean value from all technical replicates (wells) in the independent experiment.

### Overexpression of mAKT1 attenuates the expression of genes related to autophagic pathways while PFFs treatment upregulates α-syn degrading enzymes

There are multiple studies demonstrating that autophagy is playing a crucial role in α-syn degradation and autophagy impairment is involved in the pathology of synucleinopathies [35]. To illuminate the mechanism of how AKT signalling eliminates α-syn aggregation, the expression of autophagy markers was assessed with qPCR (Figure 4A-D). Our data show that overexpression of mAKT1 decreased the expression of an autophagy activator Beclin1 by almost 50% when compared to LV-GFP treated control cells (Figure 4A) regardless of PFFs treatment (p<0.05 for treatment effect in two-way RM ANOVA). Interestingly, mAKT1 overexpression also, to some extent, downregulated the expression of other autophagy-related genes, i.e. Lysosomal-associated membrane protein 1 (Lamp1) (Figure 4B) and Transcription Factor EB (TFEB) (Figure 4C). However, these changes were small and might not be related to mAKT1 action since similar effects were observed after GFP overexpression in the transduction control group. Importantly, PFFs treatment upregulated the expression of enzymes linked with the degradation of α-syn, namely Insulin Degrading Enzyme (IDE) (Figure 4E), Cathepsin B (Figure 4F) and Cathepsin D (Figure 4G). Furthermore, there was a small tendency for an increase in IDE and cathepsins expression after mAKT1 overexpression; however, this reached statistical significance only for IDE in PBS treated group (Figure 4E).

**Figure 4.**
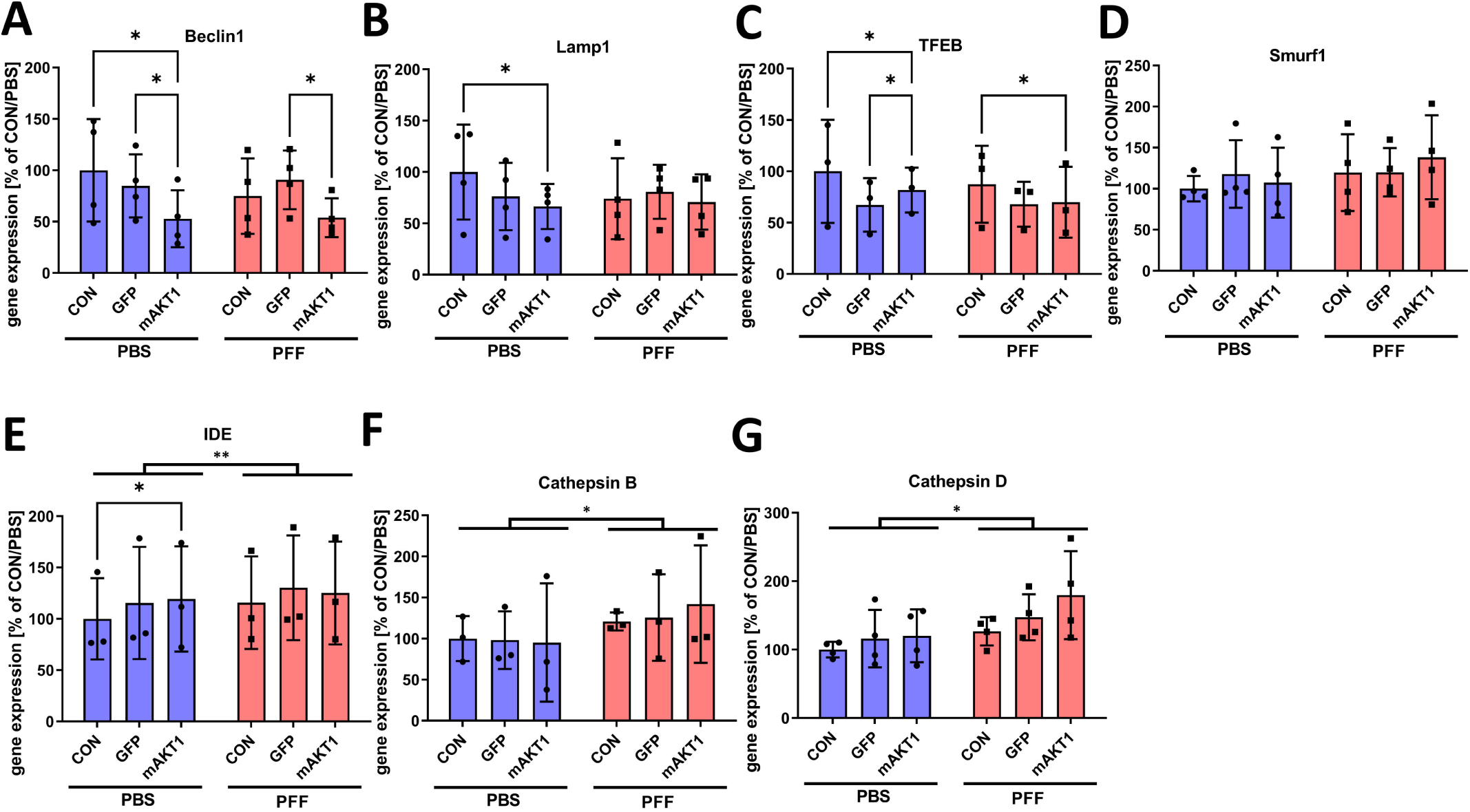
Analysis of autophagy pathway-related gene expression in hippocampal primary neurons, treated with LVs (GFP and mAKT1) and PFFs. mAKT1 attenuates expression of autophagic pathway genes: A) Beclin1, B) Lamp1, C) TFEB, but not D) Smurf1. PFFs treatment induces expression of α-syn degrading enzymes: E) Insulin Degrading Enzyme (IDE), F) Cathepsin B, G) Cathepsin D. ^*^—p-value < 0.05, ^**^—p-value < 0.01. Data are represented as mean ± SD. Each dot represents a mean value from all technical replicates (wells) in the independent experiment.

### Cathepsins B and D are essential for the protective effect of mAKT1 against α-syn aggregation

We decided to further investigate the involvement of protein degradation pathways and specific proteases in the observed effect of mAKT1 (Figure 5). First, we tested general inhibitors of autophagy (Bafilomycin A) (Figures S2B and C) and proteasome activity (Lactacystin) (Figures S2D and E) for 72 hours. Treatments with smaller concentrations of these drugs, which did not cause significant cell loss, were unable to block the protective effect of mAKT1 against α-syn aggregation, neither they exacerbated α-syn pathology in the LV-GFP control group (Figure S2B and D). Higher concentrations were also tested, but longer incubation times resulted in high toxicity that made results inconclusive (Figure S2C and E). Therefore, we decided to block specific enzymes linked to the degradation of α-syn: Cathepsins B, D and L [15,16,36,37]. Furthermore, our qPCR data show that PFFs induced upregulation of Cathepsins B and D mRNAs (Figures 4F and 4G).

**Figure 5.**
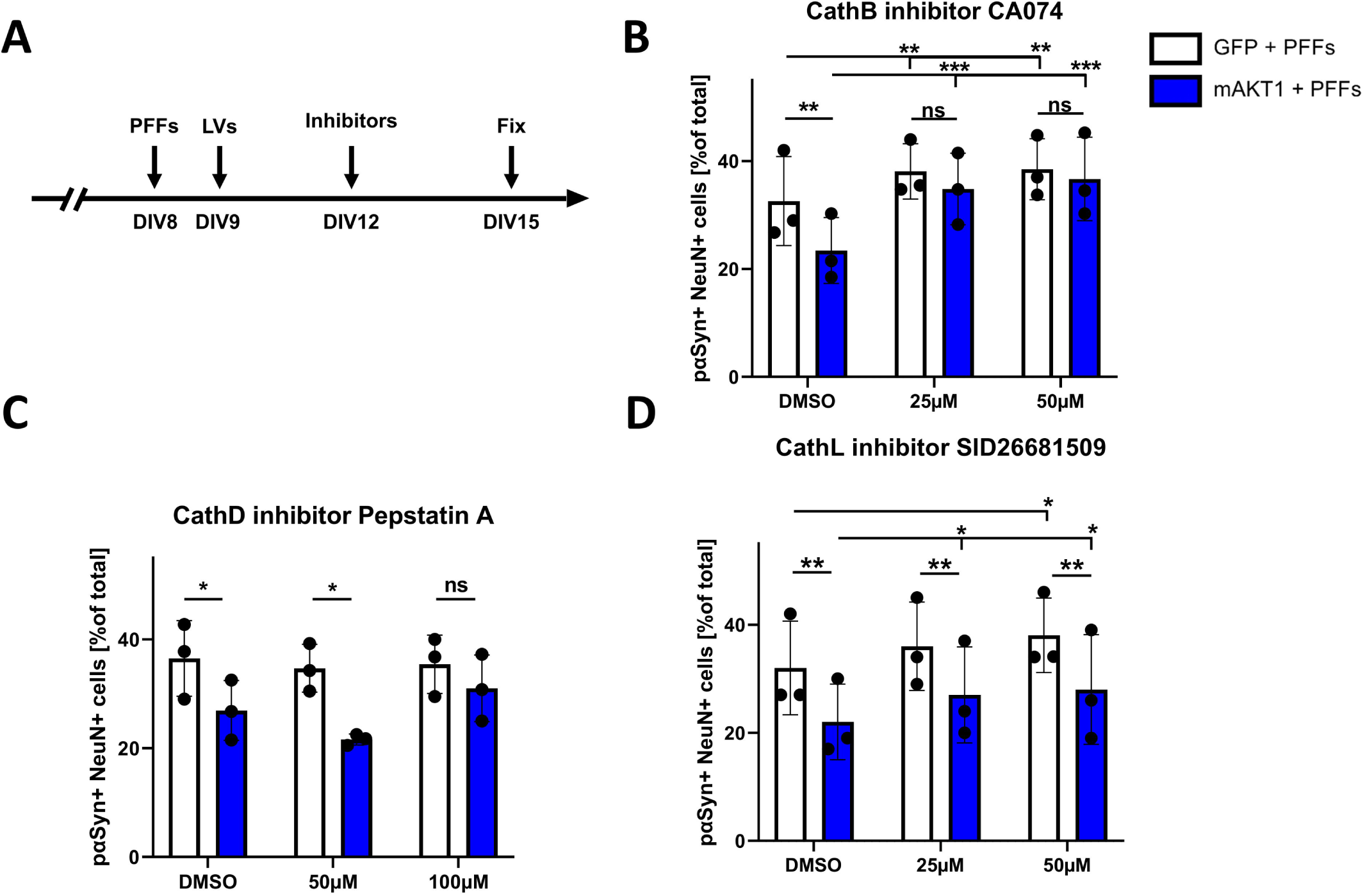
Cathepsin B and D, but not L, are essential for a protective effect of mAKT1 against α-syn aggregation. A) Treatment timeline. B) Treatment with Cathepsin B inhibitor CA074 abolished mAKT1 ability to reduce the number of Ser129-phosphorylated α-syn (pαSyn) aggregates in hippocampal neurons (NeuN+). C) Cathepsin D inhibitor Pepstatin A attenuates mAKT1 effect on the α-syn accumulation in hippocampal neurons. D) Cathepsin L inhibitor SID26681509 treatment has no effect on the ability of mAKT1 expression to alleviate α-syn aggregation in primary hippocampal neurons. ns – not significant, ^*^—p-value < 0.05, ^**^—p-value < 0.01. Data are represented as mean ± SD. Each dot represents a mean value from all technical replicates (wells) in the independent experiment.

To test the role of cathepsins in the protective effect of mAKT1 against α-syn aggregation, neurons were treated with inhibitors CA074 (Cathepsin B), Pepstatin A (Cathepsin D) or SID26681509 (Cathepsin L) (Figure 5). In this experiment, the cells were transduced at DIV9 and treated with cathepsin inhibitors at DIV12 at the time when mAKT is fully expressed. This relatively late time point treatment might have attenuated the effectiveness of both mAKT and cathepsin inhibitors; however, it allowed us to limit the potential toxic effect of compounds by reducing treatment time to 72h. DIV12 is the latest time point when in our previous studies, neurotrophic stimulation was able to reduce α-syn aggregation at full effect [12]. Moreover, inhibition of the AKT pathway on DIV12 was sufficient to increase α-syn aggregation. We speculate, based on literature data on the fate of uptaken α-syn PFFs [19], that at DIV12 α-syn PFFs uptaken by cells are still in the endolysosomal pathway, thus amenable to treatments influencing lysosomal function, while at later time points they escape to the cytoplasm.

CA074 abolished the effect of mAKT1 on α-syn accumulation (Figure 5B), as mAKT1 was effective only in DMSO-treated groups in this experiment. Main effect of mAKT1 (p<0.05) and mAKT1 × CA074 interaction (p<0.05) were significant. Interestingly, there was also a significant main effect of CA074 alone (p<0.01), indicating that in LV-GFP treated cells, Cathepsin B also plays a role in protection against α-syn aggregation, albeit the effect was small. Pepstatin A effects were less clear, with only a significant main effect of mAKT1, which was effective in DMSO and 50 μM Pepstatin A group; however, mAKT1 protective effect was abolished to some extent in 100 μM Pepstatin A group (Figure 5C). Treatment with Pepstatin A alone has not affected the accumulation of α-syn aggregates. Lastly, SID26681509 failed to inhibit mAKT1 action in both tested concentrations (Figure 5D). Again, however, apart from the clear main effect of mAKT1 (p<0.01), SID26681509 alone seemed to exacerbate α-syn aggregation as evident by the significant main effect of this inhibitor (p<0.05), albeit when analysed separately effect reached statistical significance only for the highest concentration (Figure 5D). There were no significant effects on neuronal survival of either CA074 or Pepstatin A in any tested concentrations. However, SID26681509 slightly reduced cell survival at the highest concentration, regardless of PFF or LV treatment (Figure S3).

### mAKT1 reduces the accumulation of α-syn aggregates in vivo

Finally, we tested if mAKT1 overexpression has similar effects on α-syn aggregation *in vivo*. We injected LV-mAKT1 or LV-GFP into the frontal cortex area of mice and subsequently, one week after LV injection, induced progressive α-syn aggregation through nigrostriatal injection of α-syn PFFs (Figure 6A). In this model, 30 days after PFFs injection, aggregated α-syn inclusions in neuronal somas can be seen in multiple brain structures, including cortical areas [12]. Compared to control cells transduced with LV-GFP, the population of LV-mAKT1 transduced cells had ∼35% less α-syn inclusions (Figures 6B and 6C), corroborating our *in vitro* data.

**Figure 6.**
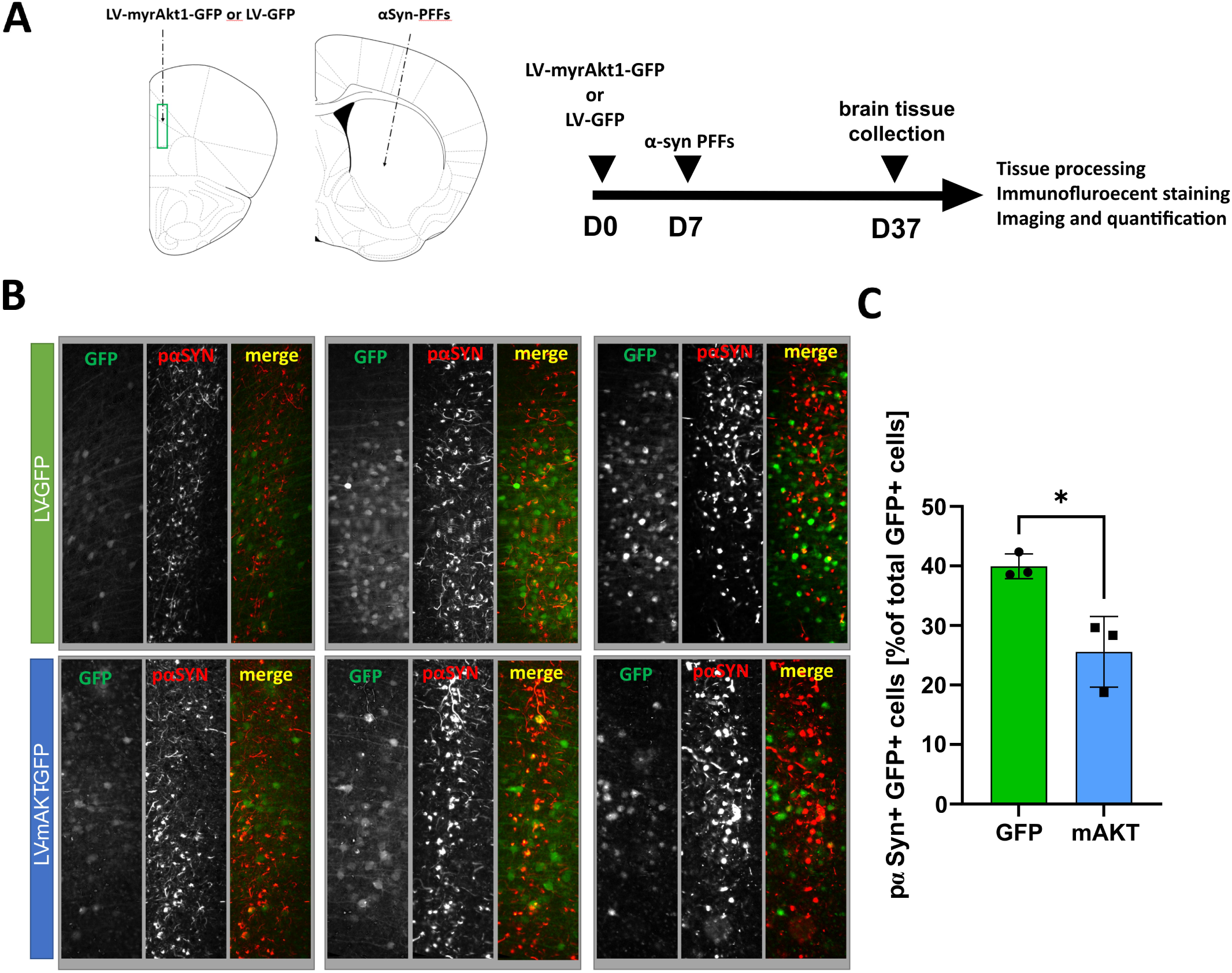
LV-delivery of mAKT1 reduces the accumulation of α-syn in cortical cells *in vivo*. A) Experiment schema: LVs were injected into the prefrontal cortex and αSyn-PFFs into the striatum as marked with dotted arrows. The green rectangle represents imaged and quantified area. Lentiviruses were injected 7 days before αSyn-PFFs. 30 days after PFFs treatment, brains were collected, stained and quantified. B) Images from the cortex of 3 animals injected with LV-GFP and 3 animals injected with LV-myrAKT1 stained for GFP (green) and pS129-αSyn (red). C) Quantification of α-syn aggregates in transduced neurons from stained cortical areas. ^*^—p-value < 0.05. Data are represented as mean ± SD, n=3 animals per group.

## Discussion

PKB/AKT1 pathway is an intracellular signalling hub activated by numerous neurotrophic factors such as glial cell line-derived neurotrophic factor (GDNF), brain-derived neurotrophic factor (BDNF), nerve growth factor (NGF), insulin-like growth factor and others that have well-known pro-survival and neurorestorative properties [38]. Not surprisingly, direct activation of PKB/AKT1 pathway has been investigated as a possible neuroprotective intervention for neurodegenerative disorders, including PD [7–11]. Our data demonstrated for the first time that constitutive activation of PKB/AKT1 pathway attenuates the accumulation of pαSyn – a histological hallmark of pathological protein aggregates found in PD and other synucleinopathies. We show that overexpression of constitutively active PKB/AKT1 – mAKT1 - can reduce α-syn burden in cultured dopamine and hippocampal neurons and confirm its effectiveness *in vivo*. Furthermore, we confirm the general pro-survival effects of LV-mediated mAKT1 overexpression in dopamine neurons and its ability to reduce ER stress-related cell death. In addition, we showed that the activity of Cathepsin B and D, but not L, is crucial for the ability of AKT1 to alleviate α-syn aggregation. Overall, our data strongly support the idea of AKT1 activation as a therapeutic strategy for the treatment of PD and other synucleinopathies.

Endosome-lysosomal and autophagy-lysosomal pathways are closely interlinked degradation pathways strongly implicated in the pathogenesis of PD. Several genes identified in familial PD are now known to be related to those pathways, i.e. LRRK2, GBA, ATP13A2 [39], while recently new risk loci for PD, components of the autophagy-lysosomal pathway, have also been reported [40,41]. There are also several lines of evidence demonstrating that aggregated α-syn can disrupt endocytic and autophagy pathways, while at the same time, those pathways are the main modes of aggregated α-syn degradation [39,42]. Moreover, endocytosis seems to be the main mode of uptake of transmittable α-syn fibrils. In fact, uptaken PFFs linger for several days in the endosomal pathway before escaping to the cytoplasm [18,19,43,44]. Internalized fibrils seem to disrupt the endocytic pathway, possibly by modulating lysosomal pH on which activity of lysosomal proteases is dependent and subsequently leads to the rupture of endocytic vesicles [42,45]. The main lysosomal proteases linked to the degradation of α-syn are Cathepsin B and D [15,46,47], while other enzymes like IDE have also been implicated [48,49]. Our data showing upregulation of Cathepsins B and D, and IDE mRNA after PFFs treatment are therefore consistent with the literature, and support the involvement of these enzymes in α-syn processing. Increases in Cathepsin B and D expression caused by mAKT1 in PFFs treated group were not significant; however, it should be noted that the activity of these enzymes is regulated on multiple posttranscriptional levels [50]. Indeed, data from experiments with cathepsins inhibitors confirm the crucial importance of Cathepsins B and D for observed effects of mAKT1 on α-syn accumulation. Interestingly, their activity is regulated by lysosomal pH, which itself can be affected by aggregated α-syn. Strikingly, it has been recently demonstrated that AKT can rescue lysosomal pH through direct interaction with lysosomal K^+^ channel TMEM175 in response to growth factors and that ablation of TMEM175 causes an increase in α-syn aggregate accumulation [24]. Taken together, we speculate that the mechanism of action by which mAKT1 protects neurons from α-syn aggregate accumulation is the normalization of endolysosomal or autophagy-lysosomal pathways function, putatively through the restoration of lysosomal pH and thus Cathepsins’ B and D activity. Interestingly, even without overexpression of mAKT1, inhibition of Cathepsin B and L, but not Cathepsin D, caused a small but significant increase in α-syn aggregate containing cells. This would suggest that the activity of these cathepsins is, to some extent protecting cells from α-syn aggregation in control PFF-treated cells. Nonetheless, the effects of observed inhibition of Cathepsins B and L were very modest, which might be due to functional redundancy between cathepsins on their action on α-syn fibrils. We were not able to utilize nonspecific inhibitors due to their toxic effects on neurons during the experiment timespan.

Further confirmation of AKT1-mediated activation of lysosomal cathepsins will require additional studies. Moreover, normalization of lysosomal pH is not the only mode by which PKB/AKT pathway can regulate autophagy – in fact, PKB/AKT is a known negative regulator of autophagy through PI3K-AKT-mTOR pathway, which is consistent with observed downregulation of Beclin1, Lamp1 and TFEB mRNA. However, considering that we only investigated these genes on mRNA level and that endosome-lysosomal and autophagy-lysosomal pathways are dysregulated by α-syn aggregation [39], our data do not allow us to infer what would be the net effect of mAKT1 on autophagy. Present data also does not allow us to discriminate if prevention of α-syn aggregation is due to increased degradation of uptaken α-syn fibrils in the endocytic pathway, degradation of α-syn aggregates forming in the cytoplasm, or both. We have, however, excluded the possibility that mAKT1 lowers endogenous α-syn levels.

We have previously demonstrated that GDNF attenuates the accumulation of α-syn aggregates both *in vitro* and *in vivo* [12] in dopamine neurons. Utilizing CRISPR/Cas9 and small molecule inhibitors, we demonstrated that the effects were dependent on RET tyrosine kinase and pointed toward the involvement of Src and AKT pathways downstream of RET. Current work supports the hypothesis that AKT activation is indeed one of the downstream mediators of GDNF effects on α-syn aggregation. Consequently, this also implies that regulating autophagy/lysosomal pathway activity is a plausible mechanism for previously reported GDNF action.

Apart from the utilization of mAKT1 as a tool for exploring the role of PKB/AKT pathway in α-syn aggregation, mAKT1 might have more direct usage in PD therapy. Upon PD diagnosis, already more than 30-50% of dopamine neurons are lost in PD patient brains while their striatal projections are even more affected with loss of about 50-80% of striatal innervation [51]. Furthermore, in just a few years of disease progression, most of the remaining cells are gone. This cell death dynamic clearly shows that neuroprotective treatments might be effective only at very early stages of the disease [38], while approaches promoting axonal regeneration or transplantation of dopamine neurons would be necessary at mid-to late-stage of the disease. Hitherto, literature data suggest that AKT activation could serve both as neuroprotective treatment [10] and promote axonal regrowth in either remaining or transplanted dopamine neurons [11]. Our data further support the therapeutic potential of AKT activation in PD – specifically, the possibility to use a constitutively active form of AKT – mAKT1. We demonstrated strong pro-survival effects of mAKT1 on cultured dopamine neurons – similar to GDNF, one of the strongest pro-survival factors for these cells. Moreover, we demonstrated the ability of mAKT1 to protect dopamine neurons from ER stress, which is linked to many neurodegenerative disorders [52,53]. Most importantly, however, we are the first to show that overexpression of constitutively active mAKT1 could protect neurons from pathological protein aggregation. Potentially this could have implications for the future development of neurorestorative and cell transplantation therapies for PD. Activating AKT with a small molecule activator is feasible, and has been demonstrated to have neuroprotective effects [54,55]. However, since AKT pathway activation is strongly implicated in cancer [56], systemic and prolonged activation of this pathway, which most probably would be necessary to achieve a disease-modifying effect on PD, rises serious safety concerns. A safer approach might be restricting AKT activation to neurons which are post-mitotic cells, e.g. by overexpression of constitutively active AKT under neuron-specific promoter. With current delivery methods, gene therapy approaches are unfeasible in early-stage PD patients, where small molecule treatments are still our best hope. However, one could imagine mAKT1 overexpression in transplanted neurons to promote both their survival and then axonal growth, in fact, this has already been suggested [11]. Importantly, our data show that overexpression of mAKT1 in neurons could have the additional benefit of protecting from ER stress and pathological protein aggregation. This is especially important since transplanted foetal mesencephalic neurons had been shown to accumulate Lewy bodies [57], demonstrating their vulnerability after transplantation. Considering the literature and our presented findings, it is tempting to speculate that overexpression of mAKT1 in transplanted neurons could be beneficial in three ways: increasing neuron survival, promoting axonal growth and lastly, protecting neurons from the ongoing pathological process – i.e. transmission of prion-like α-syn. Nonetheless, the application of mAKT1 overexpression for neurorestorative or cell transplantation therapy will require further studies and direct confirmation of such possibility; however, our work strongly supports exploring such a strategy.

## Acknowledgements

We thank Prof. Mart Saarma for his support, critical comments and suggestions during the implementation of this project. We also thank Congjun Zheng for establishing and culturing neuronal cells, Vera Kovaleva for assistance with WB experiments and Paula Collin for help with mRNA isolation. Imaging was performed at the Light Microscopy Unit, Institute of Biotechnology, supported by HiLIFE and Biocenter Finland and at the Laboratory of in vivo and in vitro Imaging at Maj Institute of Pharmacology. This work was supported by grants from the Academy of Finland #293392, #319195 and Päivikki and Sakari Sohlberg Foundation (to A.D.); National Science Centre, Poland, grant number 2019/35/D/NZ7/03200 (Sonata 15) (to P.C.); Sigrid Juselius Foundation (to M.A.), Suomen Parkinsonsäätiö, Suomen Kulttuurirahasto (00210558 Central Fund) and Doctoral School in Health Sciences (University of Helsinki) (to J.K.); and Doctoral Programme in Drug Research, the University of Helsinki (to S.E.).

## Bibliography

1. Goedert, M.; Jakes, R.; Spillantini, M.G. The Synucleinopathies: Twenty Years On. J Parkinsons Dis 2017, 7, S51–S69, doi:10.3233/JPD-179005.

2. Braak, H.; Tredici, K.D.; Rüb, U.; de Vos, R.A.I.; Jansen Steur, E.N.H.; Braak, E. Staging of Brain Pathology Related to Sporadic Parkinson’s Disease. Neurobiology o f Aging 2003, 24, 197–211, doi:10.1016/S0197-4580(02)00065-9.

3. Anderson, J.P.; Walker, D.E.; Goldstein, J.M.; de Laat, R.; Banducci, K.; Caccavello, R.J.; Barbour, R.; Huang, J.; Kling, K.; Lee, M.; et al. Phosphorylation of Ser-129 Is the Dominant Pathological Modification of Alpha-Synuclein in Familial and Sporadic Lewy Body Disease. J B iol Chem 2006, 281, 29739–29752, doi:10.1074/jbc.M600933200.

4. Borghammer, P. The α-Synuclein Origin and Connectome Model (SOC Model) of Parkinson’s Disease: Explaining Motor Asymmetry, Non-Motor Phenotypes, and Cognitive Decline. J Parkinsons Dis 2021, 11, 455–474, doi:10.3233/JPD-202481.

5. Wong, Y.C.; Krainc, D. α-Synuclein Toxicity in Neurodegeneration: Mechanism and Therapeutic Strategies. Nat Med 2017, 23, 1–13, doi:10.1038/nm.4269.

6. Mahul-Mellier, A.-L.; Burtscher, J.; Maharjan, N.; Weerens, L.; Croisier, M.; Kuttler, F.; Leleu, M.; Knott, G.W.; Lashuel, H.A. The Process of Lewy Body Formation, Rather than Simply α-Synuclein Fibrillization, Is One of the Major Drivers of Neurodegeneration. Proceedings of the National Academy of Sciences 2020, 117, 4971–4982, doi:10.1073/pnas.1913904117.

7. Diaz-Ruiz, O.; Zapata, A.; Shan, L.; Zhang, Y.; Tomac, A.C.; Malik, N.; de la Cruz, F.; Bäckman, C.M. Selective Deletion of PTEN in Dopamine Neurons Leads to Trophic Effects and Adaptation of Striatal Medium Spiny Projecting Neurons. PLoS One 2009, 4, e7027, doi:10.1371/journal.pone.0007027.

8. Domanskyi, A.; Geissler, C.; Vinnikov, I.A.; Alter, H.; Schober, A.; Vogt, M.A.; Gass, P.; Parlato, R.; Schütz, G. Pten Ablation in Adult Dopaminergic Neurons Is Neuroprotective in Parkinson’s Disease Models. FASEB J 2011, 25, 2898–2910, doi:10.1096/fj.11-181958.

9. Pellman, D.; Garber, E.A.; Cross, F.R.; Hanafusa, H. An N-Terminal Peptide from P60src Can Direct Myristylation and Plasma Membrane Localization When Fused to Heterologous Proteins. Nature 1985, 314, 374–377, doi:10.1038/314374a0.

10. Ries, V.; Henchcliffe, C.; Kareva, T.; Rzhetskaya, M.; Bland, R.; During, M.J.; Kholodilov, N.; Burke, R.E. Oncoprotein Akt/PKB Induces Trophic Effects in Murine Models of Parkinson’s Disease. Proc Natl Acad Sci U S A 2006, 103, 18757–18762, doi:10.1073/pnas.0606401103.

11. Padmanabhan, S.; Burke, R.E. Induction of Axon Growth in the Adult Brain: A New Approach to Restoration in Parkinson’s Disease. Mov Disord 2018, 33, 62–70, doi:10.1002/mds.27209.

12. Chmielarz, P.; Er, Ş.; Konovalova, J.; Bandres, L.; Hlushchuk, I.; Albert, K.; Panhelainen, A.; Luk, K.; Airavaara, M.; Domanskyi, A. GDNF/RET Signaling Pathway Activation Eliminates Lewy Body Pathology in Midbrain Dopamine Neurons. Mov Disord 2020, 35, 2279–2289, doi:10.1002/mds.28258.

13. Stefanis, L.; Emmanouilidou, E.; Pantazopoulou, M.; Kirik, D.; Vekrellis, K.; Tofaris, G.K. How Is Alpha-Synuclein Cleared from the Cell? J Neurochem 2019, 150, 577–590, doi:10.1111/jnc.14704.

14. Cullen, V.; Lindfors, M.; Ng, J.; Paetau, A.; Swinton, E.; Kolodziej, P.; Boston, H.; Saftig, P.; Woulfe, J.; Feany, M.B.; et al. Cathepsin D Expression Level Affects Alpha-Synuclein Processing, Aggregation, and Toxicity in Vivo. Mol Brain 2009, 2, 5, doi:10.1186/1756-6606-2-5.

15. McGlinchey, R.P.; Lee, J.C. Cysteine Cathepsins Are Essential in Lysosomal Degradation of α-Synuclein. Proc Natl Acad Sci U S A 2015, 112, 9322–9327, doi:10.1073/pnas.1500937112.

16. Qiao, L.; Hamamichi, S.; Caldwell, K.A.; Caldwell, G.A.; Yacoubian, T.A.; Wilson, S.; Xie, Z.-L.; Speake, L.D.; Parks, R.; Crabtree, D.; et al. Lysosomal Enzyme Cathepsin D Protects against Alpha-Synuclein Aggregation and Toxicity. Mol Brain 2008, 1, 17, doi:10.1186/1756-6606-1-17.

17. Sevlever, D.; Jiang, P.; Yen, S.-H.C. Cathepsin D Is the Main Lysosomal Enzyme Involved in the Degradation of Alpha-Synuclein and Generation of Its Carboxy-Terminally Truncated Species. Biochemistry 2008, 47, 9678–9687, doi:10.1021/bi800699v.

18. Bieri, G.; Gitler, A.D.; Brahic, M. Internalization, Axonal Transport and Release of Fibrillar Forms of Alpha-Synuclein. Neurobiology o f Disease 2018, 109, 219–225, doi:10.1016/j.nbd.2017.03.007.

19. Karpowicz, R.J.; Haney, C.M.; Mihaila, T.S.; Sandler, R.M.; Petersson, E.J.; Lee, V.M.-Y. Selective Imaging of Internalized Proteopathic α-Synuclein Seeds in Primary Neurons Reveals Mechanistic Insight into Transmission of Synucleinopathies. J B iol Chem 2017, 292, 13482– 13497, doi:10.1074/jbc.M117.780296.

20. Apetri, M.M.; Harkes, R.; Subramaniam, V.; Canters, G.W.; Schmidt, T.; Aartsma, T.J. Direct Observation of α-Synuclein Amyloid Aggregates in Endocytic Vesicles of Neuroblastoma Cells. PLoS One 2016, 11, e0153020, doi:10.1371/journal.pone.0153020.

21. Billingsley, K.J.; Bandres-Ciga, S.; Saez-Atienzar, S.; Singleton, A.B. Genetic Risk Factors in Parkinson’s Disease. Cell Tissue Res 2018, 373, 9–20, doi:10.1007/s00441-018-2817-y.

22. Perrett, R.M.; Alexopoulou, Z.; Tofaris, G.K. The Endosomal Pathway in Parkinson’s Disease. Mol C ell Neurosci 2015, 66, 21–28, doi:10.1016/j.mcn.2015.02.009.

23. Krohn, L.; Öztürk, T.N.; Vanderperre, B.; Ouled Amar Bencheikh, B.; Ruskey, J.A.; Laurent, S.B.; Spiegelman, D.; Postuma, R.B.; Arnulf, I.; Hu, M.T.M.; et al. Genetic, Structural, and Functional Evidence Link TMEM175 to Synucleinopathies. Ann Neurol 2020, 87, 139–153, doi:10.1002/ana.25629.

24. Wie, J.; Liu, Z.; Song, H.; Tropea, T.F.; Yang, L.; Wang, H.; Liang, Y.; Cang, C.; Aranda, K.; Lohmann, J.; et al. A Growth-Factor-Activated Lysosomal K+ Channel Regulates Parkinson’s Pathology. Nature 2021, 591, 431–437, doi:10.1038/s41586-021-03185-z.

25. Er, S.; Hlushchuk, I.; Airavaara, M.; Chmielarz, P.; Domanskyi, A. Studying Pre-Formed Fibril Induced α-Synuclein Accumulation in Primary Embryonic Mouse Midbrain Dopamine Neurons. J V is Exp 2020, doi:10.3791/61118.

26. Volpicelli-Daley, L.A.; Luk, K.C.; Patel, T.P.; Tanik, S.A.; Riddle, D.M.; Stieber, A.; Meaney, D.F.; Trojanowski, J.Q.; Lee, V.M.-Y. Exogenous α-Synuclein Fibrils Induce Lewy Body Pathology Leading to Synaptic Dysfunction and Neuron Death. Neuron 2011, 72, 57–71, doi:10.1016/j.neuron.2011.08.033.

27. Planken, A.; Porokuokka, L.L.; Hänninen, A.-L.; Tuominen, R.K.; Andressoo, J.-O. Medium-Throughput Computer Aided Micro-Island Method to Assay Embryonic Dopaminergic Neuron Cultures in Vitro. J Neurosci Methods 2010, 194, 122–131, doi:10.1016/j.jneumeth.2010.10.005.

28. Seibenhener, M.L.; Wooten, M.W. Isolation and Culture of Hippocampal Neurons from Prenatal Mice. J V is Exp 2012, 3634, doi:10.3791/3634.

29. Jones, T.R.; Carpenter, A.E.; Lamprecht, M.R.; Moffat, J.; Silver, S.J.; Grenier, J.K.; Castoreno, A.B.; Eggert, U.S.; Root, D.E.; Golland, P.; et al. Scoring Diverse Cellular Morphologies in Image-Based Screens with Iterative Feedback and Machine Learning. Proc Natl Acad Sci U S A 2009, 106, 1826–1831, doi:10.1073/pnas.0808843106.

30. Kamentsky, L.; Jones, T.R.; Fraser, A.; Bray, M.-A.; Logan, D.J.; Madden, K.L.; Ljosa, V.; Rueden, C.; Eliceiri, K.W.; Carpenter, A.E. Improved Structure, Function and Compatibility for CellProfiler: Modular High-Throughput Image Analysis Software. Bioinformatics 2011, 27, 1179–1180, doi:10.1093/bioinformatics/btr095.

31. Volpicelli-Daley, L.A.; Luk, K.C.; Lee, V.M.-Y. Addition of Exogenous α-Synuclein Preformed Fibrils to Primary Neuronal Cultures to Seed Recruitment of Endogenous α-Synuclein to Lewy Body and Lewy Neurite-like Aggregates. Nat Protoc 2014, 9, 2135–2146, doi:10.1038/nprot.2014.143.

32. Polinski, N.K.; Volpicelli-Daley, L.A.; Sortwell, C.E.; Luk, K.C.; Cremades, N.; Gottler, L.M.; Froula, J.; Duffy, M.F.; Lee, V.M.Y.; Martinez, T.N.; et al. Best Practices for Generating and Using Alpha-Synuclein Pre-Formed Fibrils to Model Parkinson’s Disease in Rodents. J P arkinsons Dis 2018, 8, 303–322, doi:10.3233/JPD-171248.

33. Inesi, G.; Sagara, Y. Specific Inhibitors of Intracellular Ca2+ Transport ATPases. J M embr Biol 1994, 141, 1–6, doi:10.1007/BF00232868.

34. Chmielarz, P.; Konovalova, J.; Najam, S.S.; Alter, H.; Piepponen, T.P.; Erfle, H.; Sonntag, K.C.; Schütz, G.; Vinnikov, I.A.; Domanskyi, A. Dicer and MicroRNAs Protect Adult Dopamine Neurons. Cell D eath Dis 2017, 8, e2813, doi:10.1038/cddis.2017.214.

35. Arotcarena, M.-L.; Teil, M.; Dehay, B. Autophagy in Synucleinopathy: The Overwhelmed and Defective Machinery. Cells 2019, 8, 565, doi:10.3390/cells8060565.

36. Luhr, K.M.; Nordström, E.K.; Löw, P.; Kristensson, K. Cathepsin B and L Are Involved in Degradation of Prions in GT1-1 Neuronal Cells. Neuroreport 2004, 15, 1663–1667, doi:10.1097/01.wnr.0000134931.81690.34.

37. Mueller-Steiner, S.; Zhou, Y.; Arai, H.; Roberson, E.D.; Sun, B.; Chen, J.; Wang, X.; Yu, G.; Esposito, L.; Mucke, L.; et al. Antiamyloidogenic and Neuroprotective Functions of Cathepsin B: Implications for Alzheimer’s Disease. Neuron 2006, 51, 703–714, doi:10.1016/j.neuron.2006.07.027.

38. Chmielarz, P.; Saarma, M. Neurotrophic Factors for Disease-Modifying Treatments of Parkinson’s Disease: Gaps between Basic Science and Clinical Studies. Pharmacol Rep 2020, 72, 1195–1217, doi:10.1007/s43440-020-00120-3.

39. Teixeira, M.; Sheta, R.; Idi, W.; Oueslati, A. Alpha-Synuclein and the Endolysosomal System in Parkinson’s Disease: Guilty by Association. Biomolecules 2021, 11, 1333, doi:10.3390/biom11091333.

40. Chang, D.; Nalls, M.A.; Hallgrímsdóttir, I.B.; Hunkapiller, J.; van der Brug, M.; Cai, F.; International Parkinson’s Disease Genomics Consortium; 23andMe Research Team; Kerchner, G.A.; Ayalon, G.; et al. A Meta-Analysis of Genome-Wide Association Studies Identifies 17 New Parkinson’s Disease Risk Loci. Nat Genet 2017, 49, 1511–1516, doi:10.1038/ng.3955.

41. Chen, Y.-P.; Gu, X.-J.; Song, W.; Hou, Y.-B.; Ou, R.-W.; Zhang, L.-Y.; Liu, K.-C.; Su, W.-M.; Cao, B.; Wei, Q.-Q.; et al. Rare Variants Analysis of Lysosomal Related Genes in Early-Onset and Familial Parkinson’s Disease in a Chinese Cohort. J Parkinsons Dis 2021, 11, 1845–1855, doi:10.3233/JPD-212658.

42. Wildburger, N.C.; Hartke, A.-S.; Schidlitzki, A.; Richter, F. Current Evidence for a Bidirectional Loop Between the Lysosome and Alpha-Synuclein Proteoforms. Front C ell Dev Biol 2020, 8, 598446, doi:10.3389/fcell.2020.598446.

43. Flavin, W.P.; Bousset, L.; Green, Z.C.; Chu, Y.; Skarpathiotis, S.; Chaney, M.J.; Kordower, J.H.; Melki, R.; Campbell, E.M. Endocytic Vesicle Rupture Is a Conserved Mechanism of Cellular Invasion by Amyloid Proteins. Acta Neuropathol 2017, 134, 629–653, doi:10.1007/s00401-017-1722-x.

44. Hijaz, B.A.; Volpicelli-Daley, L.A. Initiation and Propagation of α-Synuclein Aggregation in the Nervous System. Mol Neurodegener 2020, 15, 19, doi:10.1186/s13024-020-00368-6.

45. Hoffmann, A.-C.; Minakaki, G.; Menges, S.; Salvi, R.; Savitskiy, S.; Kazman, A.; Vicente Miranda, H.; Mielenz, D.; Klucken, J.; Winkler, J.; et al. Extracellular Aggregated Alpha Synuclein Primarily Triggers Lysosomal Dysfunction in Neural Cells Prevented by Trehalose. Sci Rep 2019, 9, 544, doi:10.1038/s41598-018-35811-8.

46. Stoka, V.; Turk, V.; Turk, B. Lysosomal Cathepsins and Their Regulation in Aging and Neurodegeneration. Ageing R es Rev 2016, 32, 22–37, doi:10.1016/j.arr.2016.04.010.

47. Tsujimura, A.; Taguchi, K.; Watanabe, Y.; Tatebe, H.; Tokuda, T.; Mizuno, T.; Tanaka, M. Lysosomal Enzyme Cathepsin B Enhances the Aggregate Forming Activity of Exogenous α-Synuclein Fibrils. Neurobiol Dis 2015, 73, 244–253, doi:10.1016/j.nbd.2014.10.011.

48. Sharma, S.K.; Chorell, E.; Steneberg, P.; Vernersson-Lindahl, E.; Edlund, H.; Wittung-Stafshede, P. Insulin-Degrading Enzyme Prevents α-Synuclein Fibril Formation in a Nonproteolytical Manner. Sci Rep 2015, 5, 12531, doi:10.1038/srep12531.

49. Sousa, L.; Guarda, M.; Meneses, M.J.; Macedo, M.P.; Vicente Miranda, H. Insulin-Degrading Enzyme: An Ally against Metabolic and Neurodegenerative Diseases. J Pathol 2021, 255, 346– 361, doi:10.1002/path.5777.

50. Tayebi, N.; Lopez, G.; Do, J.; Sidransky, E. Pro-Cathepsin D, Prosaposin, and Progranulin: Lysosomal Networks in Parkinsonism. Trends M ol Med 2020, 26, 913–923, doi:10.1016/j.molmed.2020.07.004.

51. Kordower, J.H.; Olanow, C.W.; Dodiya, H.B.; Chu, Y.; Beach, T.G.; Adler, C.H.; Halliday, G.M.; Bartus, R.T. Disease Duration and the Integrity of the Nigrostriatal System in Parkinson’s Disease. Brain 2013, 136, 2419–2431, doi:10.1093/brain/awt192.

52. Hetz, C.; Saxena, S. ER Stress and the Unfolded Protein Response in Neurodegeneration. Nat R ev Neurol 2017, 13, 477–491, doi:10.1038/nrneurol.2017.99.

53. Kovaleva, V.; Saarma, M. Endoplasmic Reticulum Stress Regulators: New Drug Targets for Parkinson’s Disease. J Parkinsons Dis 2021, 11, S219–S228, doi:10.3233/JPD-212673.

54. Zhang, D.; Zhang, H.; Hao, S.; Yan, H.; Zhang, Z.; Hu, Y.; Zhuang, Z.; Li, W.; Zhou, M.; Li, K.; et al. Akt Specific Activator SC79 Protects against Early Brain Injury Following Subarachnoid Hemorrhage. ACS C hem Neurosci 2016, 7, 710–718, doi:10.1021/acschemneuro.5b00306.

55. Jo, H.; Mondal, S.; Tan, D.; Nagata, E.; Takizawa, S.; Sharma, A.K.; Hou, Q.; Shanmugasundaram, K.; Prasad, A.; Tung, J.K.; et al. Small Molecule-Induced Cytosolic Activation of Protein Kinase Akt Rescues Ischemia-Elicited Neuronal Death. Proc Natl Acad Sci U S A 2012, 109, 10581–10586, doi:10.1073/pnas.1202810109.

56. Alzahrani, A.S. PI3K/Akt/MTOR Inhibitors in Cancer: At the Bench and Bedside. Semin Cancer Biol 2019, 59, 125–132, doi:10.1016/j.semcancer.2019.07.009.

57. Kordower, J.H.; Chu, Y.; Hauser, R.A.; Freeman, T.B.; Olanow, C.W. Lewy Body-like Pathology in Long-Term Embryonic Nigral Transplants in Parkinson’s Disease. Nat Med 2008, 14, 504– 506, doi:10.1038/nm1747.

