## supplementary figures for "PKB/AKT attenuates Lewy Body-like pathology in primary neurons via Cathepsins B and D"

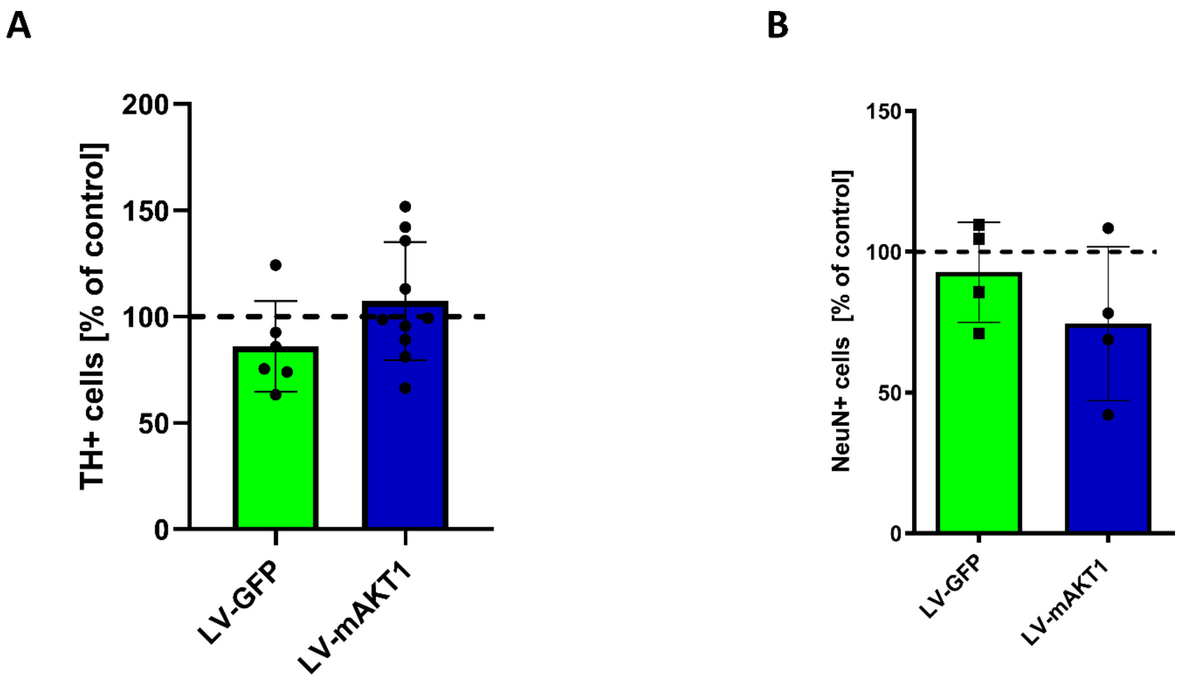


Figure S1. LV-GFP and LV-mAKT1 treatments had no significant effect on survival of primary A) midbrain and B) hippocampal neurons, when added at DIV5. Data are represented as mean ± SD. Each dot represents a mean value from all technical replicates (wells) in the independent experiment.

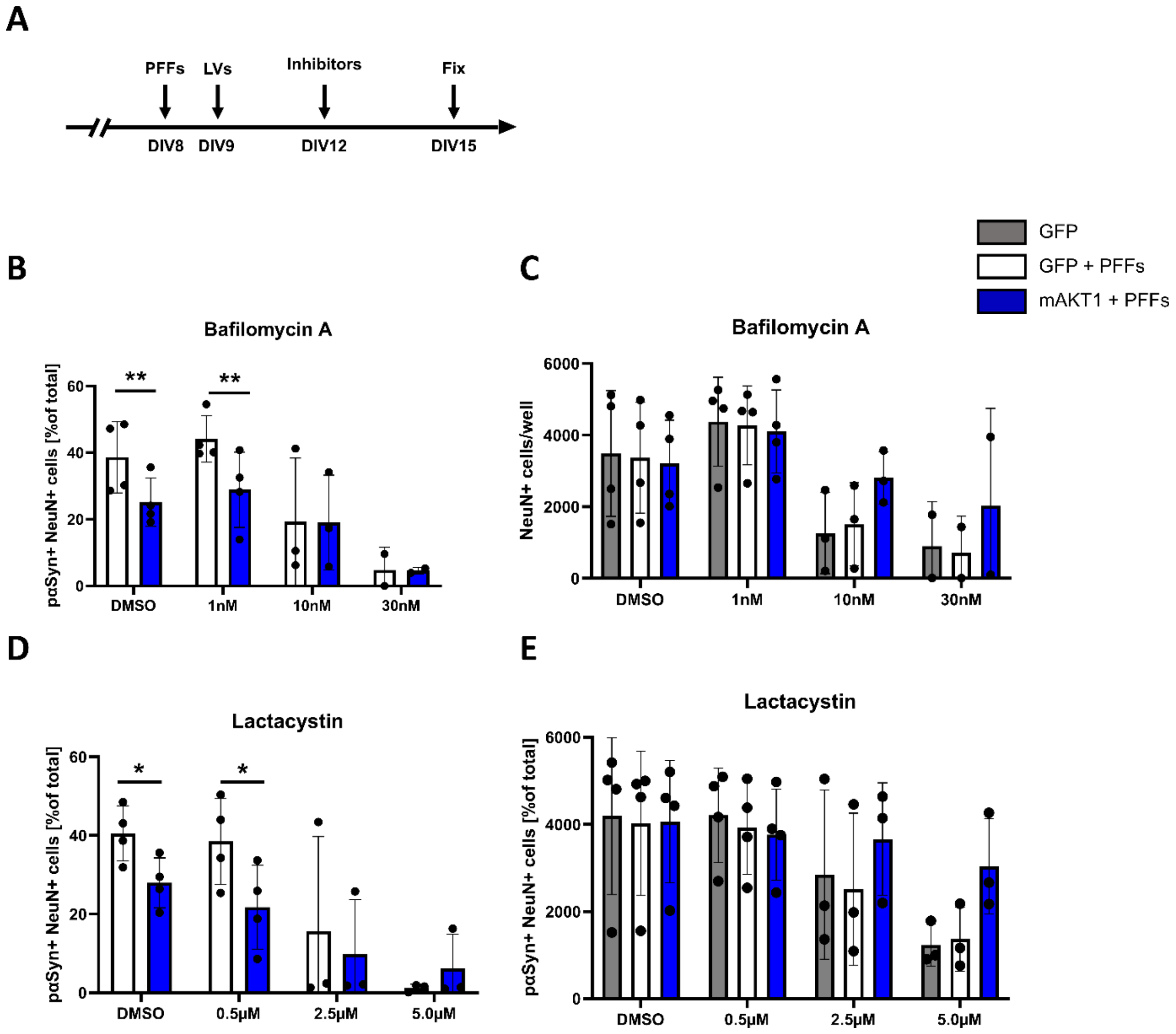


Figure S2. Treatment of hippocampal neuronal cultures with low doses of general inhibitors of autophagy (Bafilomycin A) or proteasomal activity (Lactacystin) had no effect on mAKT1 ability to reduce α-syn accumulation, while higher doses induced prominent cell toxicity. A) Treatment timeline. B) Lower dose of autophagy inhibitor Bafilomycin A did not interfere with mAKT1 ability to reduce α-syn aggregation in hippocampal primary neurons (NeuN+). C) Effect of different doses of Bafilomycin A on survival of hippocampal primary neurons. D) Treatment of hippocampal neurons with proteasome inhibitor Lactacystin did not abolish ability of mAKT1 to alleviate α-syn accumulation. E) Survival rate of hippocampal primary neurons after treatment with different doses of Lactacystin. *—p-value < 0.05, **—p-value < 0.01. Data are represented as mean ± SD. Each dot represents a mean value from all technical replicates (wells) in the independent experiment.

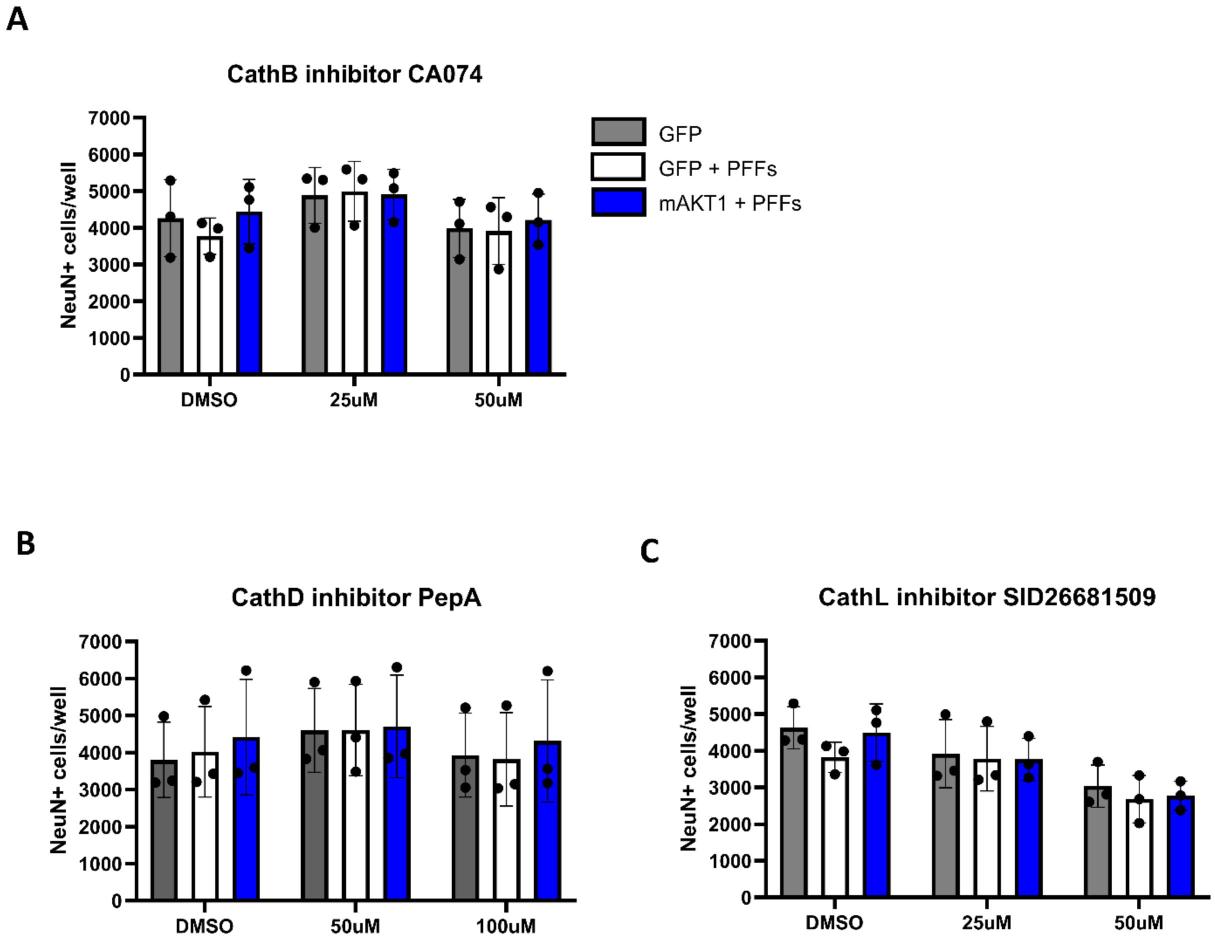


Figure S3. Survival of primary hippocampal neurons, treated with A) Cathepsin B inhibitor CA074, B) Cathepsin D inhibitor Pepstatin A, and C) Cathepsin L inhibitor SID26681509. Data are represented as mean ± SD. Each dot represents a mean value from all technical replicates (wells) in the independent experiment.
